# Design principles for open source bioinstrumentation: the poseidon syringe pump system as an example

**DOI:** 10.1101/521096

**Authors:** A. Sina Booeshaghi, Eduardo da Veiga Beltrame, Dylan Bannon, Jase Gehring, Lior Pachter

## Abstract

The poseidon syringe pump and microscope system is an open source alternative to commercial systems. It costs less than $400 and can be assembled in under an hour using the instructions and source files available at https://pachterlab.github.io/poseidon. We describe the poseidon system and use it to illustrate design principles that can facilitate the adoption and development of open source bioinstruments. The principles are functionality, robustness, simplicity, modularity, benchmarking, and documentation.

## Introduction

Open source hardware projects^1^ have become increasingly popular in recent years due in part to i) the development of an ecosystem of open source electronics boards like the Arduino and Raspberry Pi systems^2, 3^ and ii) the rapid evolution of desktop 3D printers. These developments have spurred growing interest in laboratory instrument open source projects^4–6^ including syringe pumps^7^, microscopes^8^, fluorescence imaging devices^9^, micro-dispensers^10^ and single-cell transcriptomics technologies^11^. While lower cost can be an important reason for development of open source hardware^12^, the ability to customize designs for specific applications gives open source projects a unique advantage over commercial solutions. In addition, expanding libraries of designs, software, and commonly used off-the-shelf parts can be shared and adapted across projects, meaning developers are never starting from scratch, even when designing a new instrument. For example, the RepRap project 3D printers borrowed heavily from standard software and hardware Computer Numeric Control (CNC) tools used in machining. As open source designs, electronics boards, software, and parts for 3D printers were continually published and improved, cheap and interchangeable open source hardware and software intended for 3D printing began to be repurposed for new bioinstruments such as liquid handlers^13^, vial handlers and food dispensers^14^, autosamplers^15, 16^, and bioprinters^17, 18^.

Our laboratory has a general interest in developing new methods for high-throughput single-cell applications such as Drop-seq^19^ and inDrops^20^ which rely on precise flow rate control to operate microfluidic devices. The unpredictable landscape of single-cell genomics technology puts a high priority on flexible hardware and software that can be adapted and re-purposed as experiments evolve. Frustrated by the inflexible software interface and functionality offered by commercial systems, and inspired by the array of do-it-yourself electronics and instrumentation projects powered by open source hardware, we sought to develop our own open source multi-syringe pump array and microscope system for low cost microfluidics experiments.

This system, which we call poseidon, is based on published open source syringe pumps^7^ and microscope microfluidics stations^11^ but introduces a number of innovations and adapts common 3D printer hardware and software to control the system. Requiring only off-the-shelf components and 3D printed parts, poseidon can be assembled in less than an hour for less than $400. While developing the poseidon system, we paid attention to the many design choices involved as well as their implications, leading to a set of open source hardware design principles specifically tailored to bioinstrumentation. Here we detail these design principles using the poseidon system as an illustrated example. We hope that these principles, and poseidon itself, will be of use to others and facilitate the development of other open source bioinstrumentation projects.

The poseidon syringe pump array and microscope system is an open source alternative to commercial systems (Fig. 1). It uses 3D printed parts and common components that can be easily purchased from multiple retailers. The pumps and microscope can be used together for microfluidics experiments, or the pumps can be connected to a computer and used independently. For scientists with tight budgets, the microscope system, which is stand-alone, is an effective solution for basic light microscopy.

**Figure 1.**
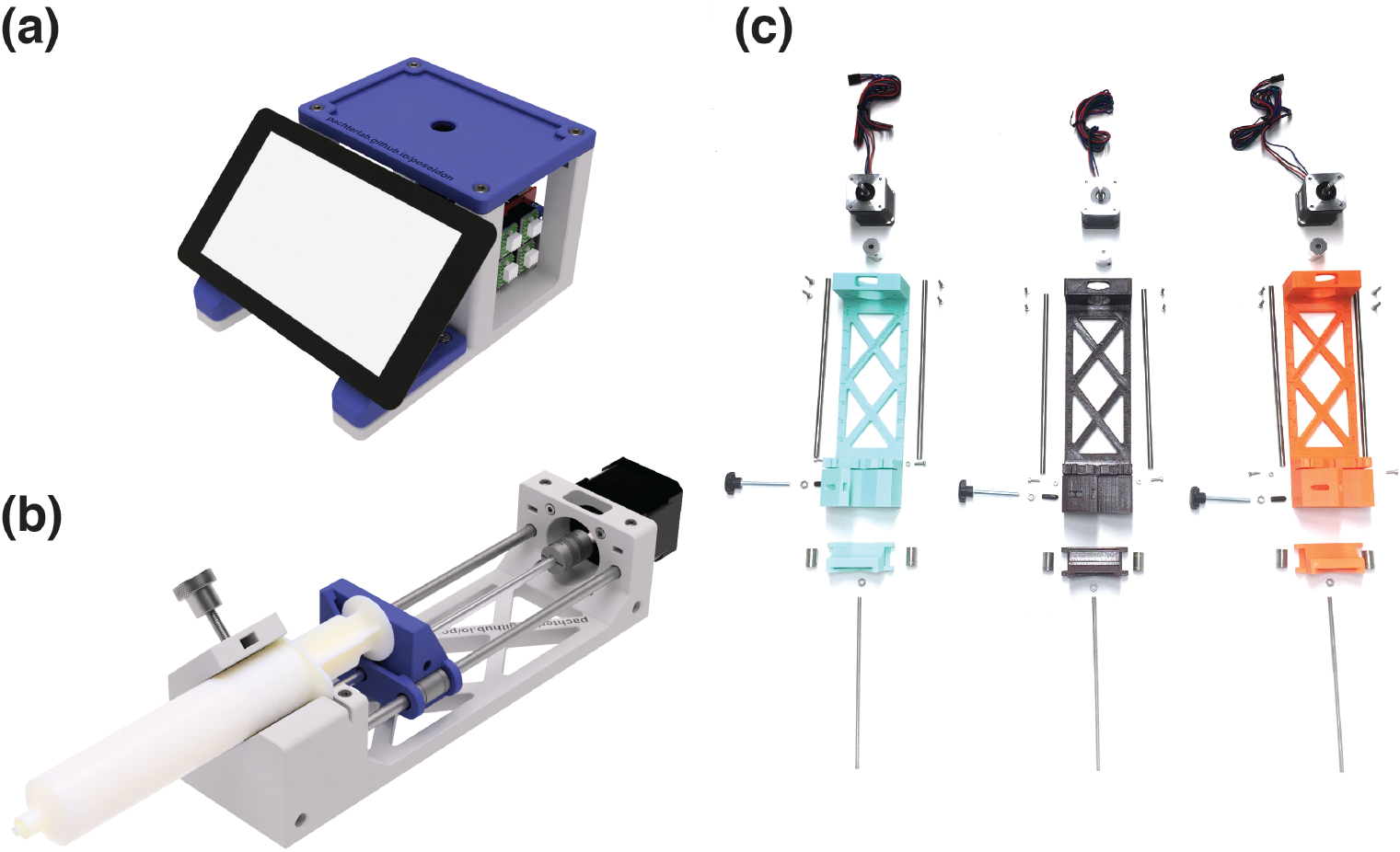
Model of the poseidon system. (a) CAD rendering of the microscope station and (b) a single syringe pump loaded with a 60 mL syringe. (c) Exploded view of all the components needed for assembling three pumps.

The poseidon system uses a Raspberry Pi and touchscreen for the microscope and an Arduino board with a CNC shield to operate up to four pumps simultaneously. Each pump has a stepper motor that drives a lead screw, which in turn moves a sled (mounted on linear bearings) that pushes (infuses) or pulls (aspirates) the syringe plunger. The microscope camera and Arduino use USB connections to connect to the Raspberry Pi or desktop computer (Fig. 2). The system was developed using readily available tools (Fig. 3).

The poseidon system repository is available under a BSD 2-clause license at https://github.com/pachterlab/poseidon. For reproducibility and ease of adoption we included direct links to the specific parts used for poseidon, in the GitHub repository. The following components are available:

1. 3D models and Computer Aided Design (CAD) files of the 3D printed components.
2. Pump controller and Graphical User Interface (GUI) software to control the Arduino.
3. Arduino firmware to relay commands via USB to drive the motors.

## Design principles

As we invested more time into poseidon, our goal became adoption by new users rather than simply meeting our own specific design requirements. This mindset shaped our entire design process. We strove to produce a system that could be readily implemented and modified by others, who could then improve the system and expand it to new use cases. We considered that bioinstrument users generally fall into two categories: i) those who want to adopt a design and use it in a straightforward manner, and ii) those who want to tweak, improve, and adapt designs to their needs and are utilizing the instrument for new use cases. While cost is one motivation for developing and using open source instruments, low cost alone cannot drive the adoption of a project for these two groups. A successful open source instrument appeals to the needs of basic and advanced users by adhering to a set of clear design principles: functionality, robustness, simplicity, modularity, benchmarking, and documentation.

### Functionality: Being good enough for the application at hand

In engineering, a functional requirement defines a specific metric that a hardware or software system must achieve. The idea of being “good enough” attempts to capture the many design decisions that can be made during the design-build-test cycle as developers consider the tradeoffs between utility, precision, accuracy, speed, cost, and complexity that are acceptable given the application. The poseidon system needed to achieve the following functional requirements for use in microfluidic applications:

1. The syringe pumps needed to be precise enough to make monodisperse emulsions on droplet generation microfluidics chips, with flow rates on the order of 1 mL/hr.
2. The microscope needed to have sufficient magnification to examine the emulsions and view the microfluidic device during operation.
3. The hardware and control software needed to be able to run at least three pumps independently.
4. The software interface needed to be simple and allow users to easily change flow rates, select syringe type or diameter, and perform gradient pumping.
5. The software needed to operate the microscope.

**Figure 2.**
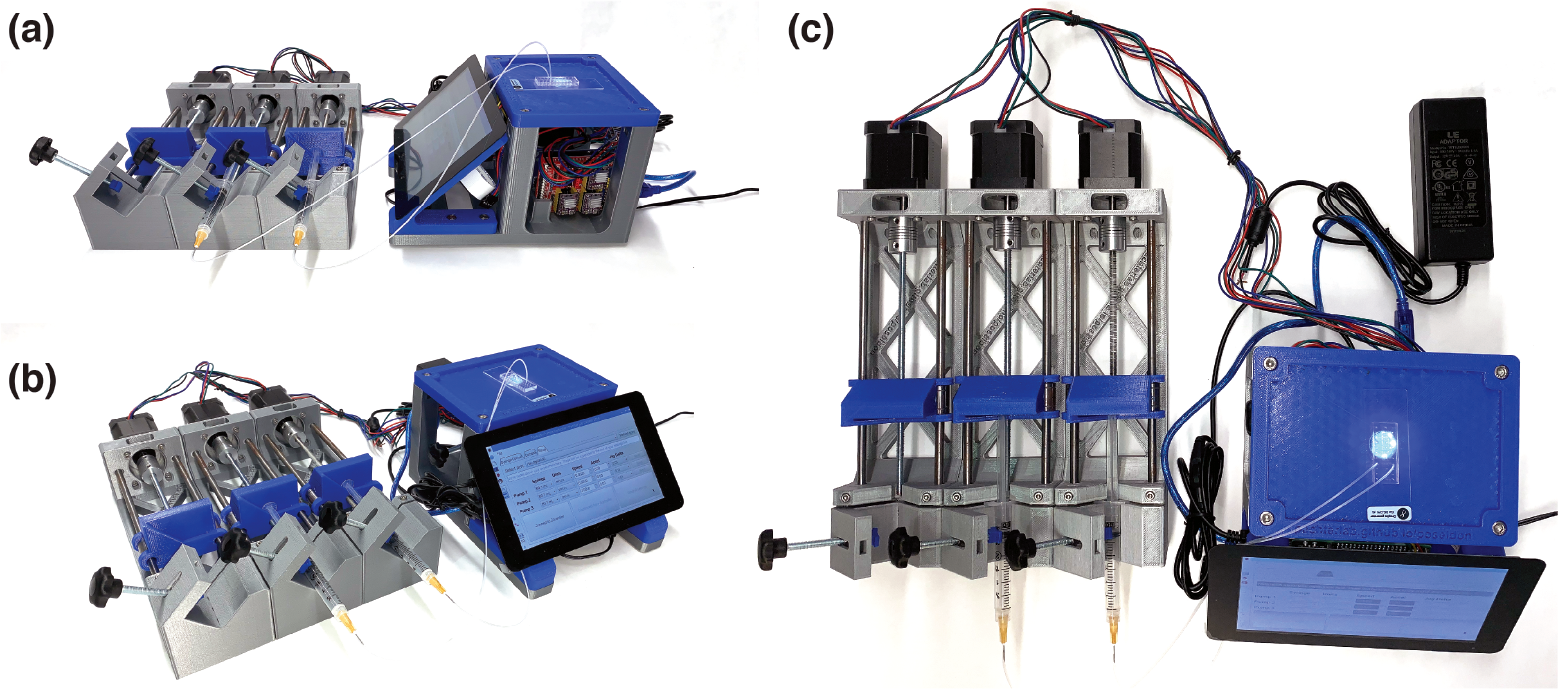
Using the poseidon system. Configuration of the poseidon system for running an emulsion generation microfluidics experiment where only two pumps are used. (a) Side view (b) Angled view (c) Top view.

These were the minimum requirements that were specified before we began developing poseidon. After designing hardware that should be able to meet these objectives, we ensured the pumps operated reliably with flow rates ranging from a few hundred microliters per hour up to several hundred milliliters per minute and we selected an inexpensive USB microscope that reliably imaged our microfluidic device.

### Robustness: Tolerating variations in manufacturing and operation

Robustness encompasses not only mitigating the possibility of failure during operation but also ensuring a construction process that tolerates variability in the components. This is particularly important in biology applications where instruments must frequently work in varying physical conditions and with variable input. Ensuring robustness took considerable time, demanding attention to small details and repeated testing. For example, much open source hardware relies on 3D printed components that can introduce variability when printed on different printers. Mechanical tolerance was built into the 3D printed parts over the course of many design-build-test cycles, such as by modifying the print settings to allow for a press fit of the syringe into the pump. During testing, we discovered an unforeseen hardware issue: when there was too much sliding resistance on the carriage, the linear rods displaced and the printed plastic body bent. To stop the bending, we designed a reinforced body and secured the linear rods with set screws.

On the software side, robustness demands testing to minimize user operation error and to ensure correct functionality. The software must be installed and tested on multiple operating systems to verify operation is as expected. Once the poseidon pumps were being used for experiments in our lab and others, usability issues became apparent. For example, one version of the software configured the stepper motors to use a different microstepping than the hardware had configured, an error which the users encountered during their experiments by observing incorrect flow rates. Using the software during an experiment also revealed small usability issues that had to be corrected, such as using a drop-down menu for choosing the flow direction instead of using a “+” or “−” sign in the displacement amount box. These improvements are relatively minor on their own, but we believe the sum total of such small modifications has an outsized impact on potential adopters testing out an unfamiliar system for the first time.

### Simplicity: Making it easy to source, build, and operate

Simplicity and ease-of-use are essential for the adoption of bioinstruments. Sourcing components for a design should be as easy as possible, prioritizing off-the-shelf components during development and incorporating harder to find parts only if necessary for the application at hand. An accurate and up-to-date bill of materials (BOM), with ideally more than one vendor for each part, simplifies purchasing and leads to easier adoption. For the poseidon system, we ensured that users would be able to purchase all the components from Amazon. During assembly, it is important to recognize that using specialized equipment - even soldering a circuit board - may be a barrier to adoption. While such specialized assembly processes are sometimes unavoidable, simplicity is paramount. An excellent way to assess the difficulty of assembly is to have people unfamiliar with the project perform the assembly using only the documentation available. With the poseidon system it was possible to design around most of these constraints, and we verified that assembly of a single pump by a new user following the instruction video takes less than 15 minutes, requiring only pliers and screwdrivers.

**Figure 3.**
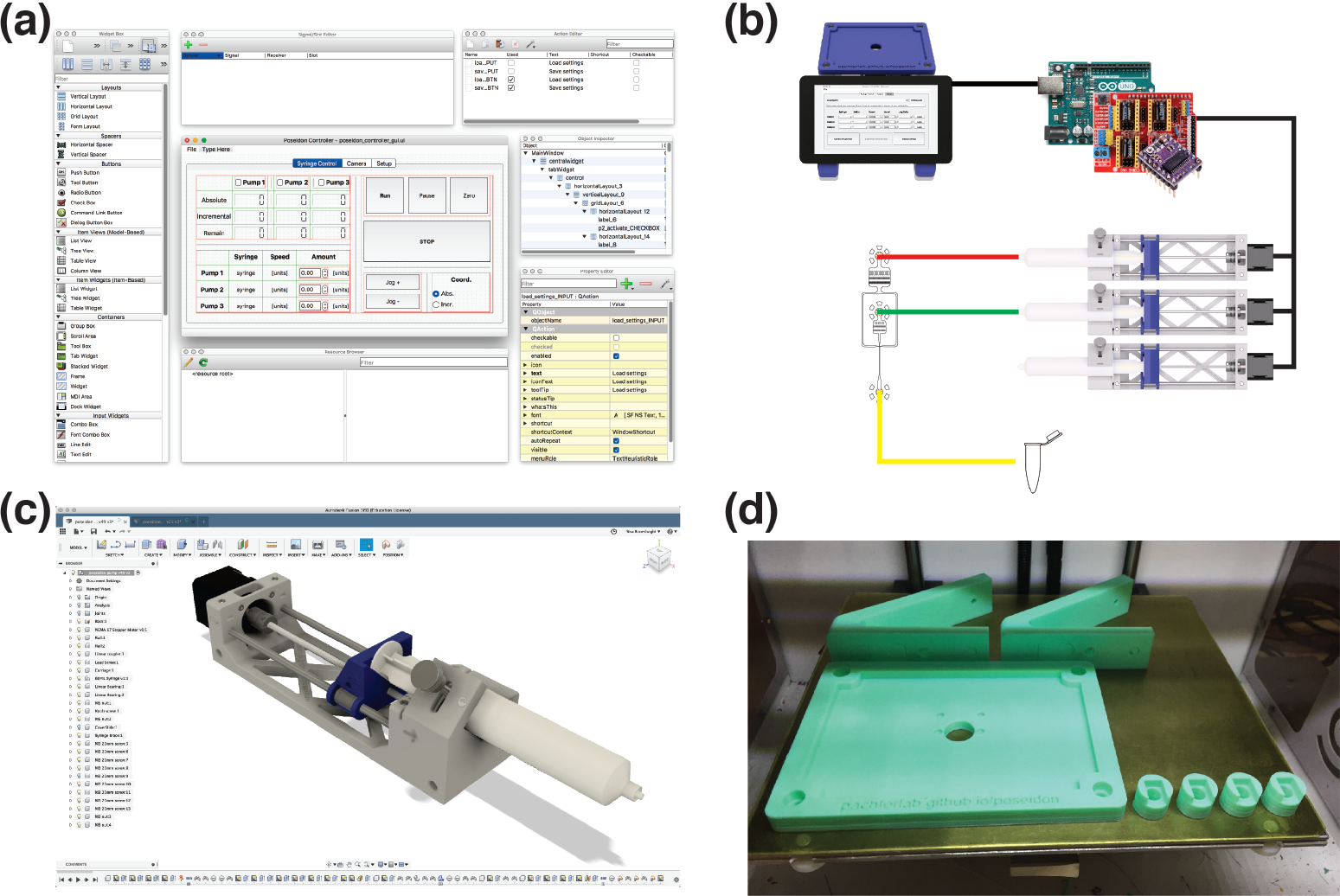
Overview of the tools used for developing the poseidon system. (a) The GUI was created using Qt Designer^21^, an open source drag and drop application, for organizing buttons that allows users to easily change the GUI interface when adding new functionalities. (b) This GUI interfaces with a Python script that controls both the microscope and Arduino via USB. The Arduino controls the stepper motors on each pump using the CNC shield and stepper motor drivers. (c) The system’s 3D printed components were designed using Fusion 360^22^, a cloud enabled CAD software that streamlines collaboration and offers free licenses for academics, hobbyists and small businesses. To modify the 3D models users can either work with Fusion 360 or any other CAD software. (d) The 3D printed components can be fabricated on any fused filament fabrication (FFF) 3D printer.

All of the above considerations also apply to software. For example, minimizing dependency on external software libraries simplifies installation and avoids versioning issues. From a user’s perspective, having a single executable file for the software is ideal. We compiled the Python scripts into single-click executable files for Mac, Windows, and Unix. The custom poseidon Arduino firmware needs be loaded onto the Arduino Uno board following simple instructions. If users wishe to use a Raspberry Pi to operate poseidon, installation requires flashing an SD card with the official version of the Raspbian OS image.

### Modularity: Designing independent and interchangeable units

Because some users will want to adapt a design to new use cases, it is important to consider how easily a design can be taken apart, tweaked, and re-purposed. A modular design with independent components that can be interfaced with each other is easier to re-purpose and improve on than a tightly integrated device. When a design is not modular, developing new features may require a complete redesign. This is the problem that led us to develop poseidon in the first place – our commercial, highly integrated system was too rigid to meet our changing needs. The software could not be improved or modified, and the small integrated touchscreen interface, marketed as an advantage^23^, was a hindrance to routine operation. In the case of poseidon, some users might want to add additional features to the pumps such as an electronic end stop. The modularity of the Arduino board makes this change straightforward and simple to implement.

A design is also easier to modify when common and standardized parts and connectors are used. Standardization is ubiquitous in both open source hardware and software projects. Open source 3D printers use a common set of screws, rods, extruding nozzles, and electronics. Often these printers are variations of a few common, popular, and proven designs. The standardization of 3D printer parts means that they, in turn, can be readily adapted for new use cases. Almost all of the components used in the poseidon system can be found in open source desktop 3D printers.

**Figure 4.**
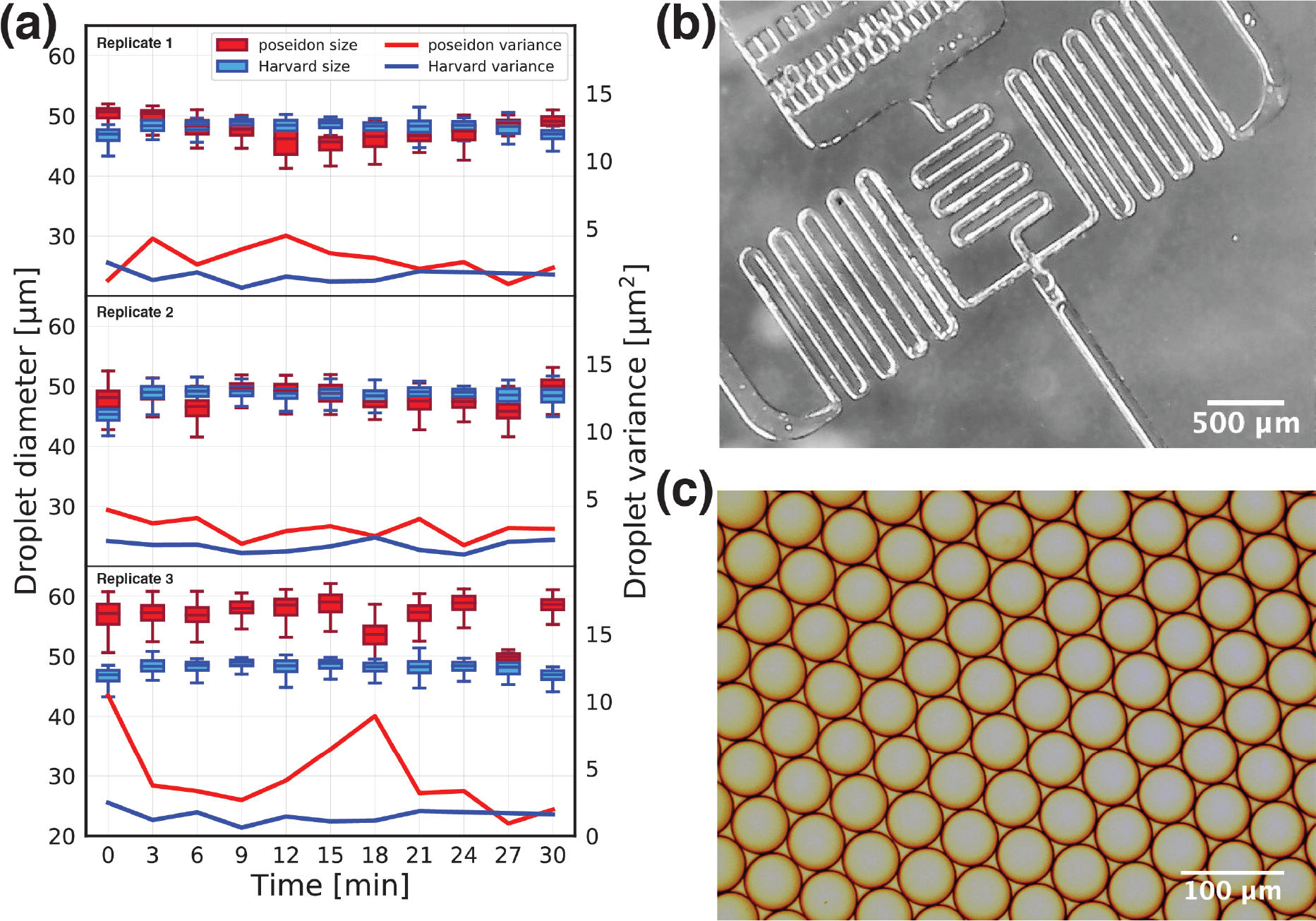
Benchmarking the poseidon system against the Harvard Apparatus system. Using a droplet generation chip we compared the droplet diameter size between two systems. (a) The poseidon system produces droplets that are close to the expected droplet size of 58 *μ*m in diameter. The variance between the droplets created with the two systems are comparable. (b) A microfluidic droplet generation chip imaged using the poseidon microscope (c) Example of a monodisperse emulsion produced by the poseidon system.

### Benchmarking: Validating with standard protocols

Users need to know the degree to which instrument design is applicable for the problem at hand. Thus it is important to describe protocols where instruments have been applied and provide benchmarking results. Open source instruments may not always perform as well commercial systems, but may still be good enough for many applications. Direct comparison with commercial instruments and clear identification of device shortcomings, or features still in development, is important to instill confidence in the system.

The poseidon system has been successfully used in generating monodisperse emulsions using the inDrops emulsion generator device (Fig. 4). We achieved targeted droplet size with small variance in droplet diameter. Importantly, we directly compared poseidon with the commercial array from Harvard Apparatus (catalog numbers 2401-408 and 70-3406), demonstrating comparable variance in droplet diameter. The poseidon system can be operated at a range of flow rates (Table 1). By physically adding or removing the microstepping jumpers on the CNC shield board users can access a range of flow rates, so that the same system can be used for high-precision microfluidics experiments and high flow rate applications such as protein purification.

### Documentation: Describing the system to others

Clear instructions and documentation are essential to facilitate rapid and painless assembly. Videos, photographs and written descriptions are fundamental for showcasing a design and ensuring adoption. For assemblies, videos are often the most helpful documentation for users and do not take much time and effort to produce. Videos also clearly convey how much effort and time users should expect to invest in assembly. Documentation enables faster and easier understanding of design.

**Figure 5.**
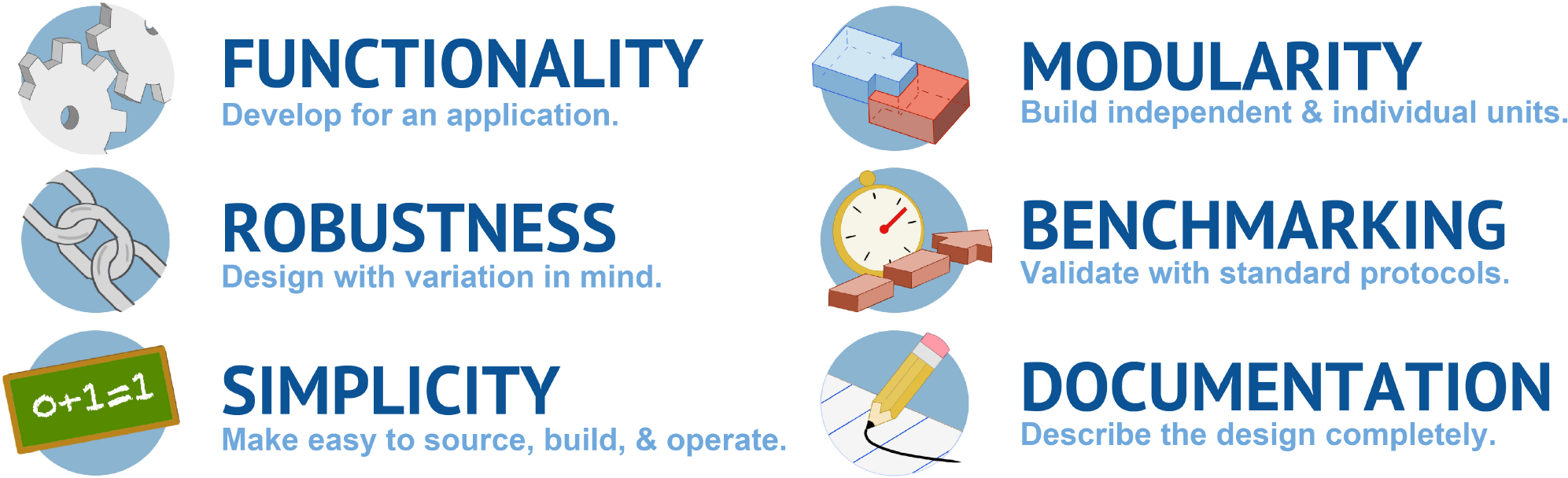
Summary of the design principles for open source bioinstrumentation.

Users who want to modify a design will additionally benefit from understanding design decisions – both those motivated by technical considerations and those motivated by user feedback. How to implement each feature is the result of thought and iteration from the designer, but what is learned may not be readily apparent in the final designs. Documentation of lessons learned, and insight into why design features were implemented a certain way is important; sometimes modifications that seem to be an improvement will create a failure mode that is not readily apparent.

For the poseidon system, multiple build videos are available on YouTube showing the entire assembly process. In making the poseidon documentation website, we also strove to use clearly labeled photos of the hardware with short written instructions, as this makes it easier for prospective users to grasp the design and expected time investment at a glance. The development and documentation of open source projects benefits tremendously from making use of version control repositories, which streamlines remote collaboration and development tracking. For the poseidon system, we used the online repository GitHub, which allows for version control and documentation of each change made and makes it simple to create user guides and documentation as can be seen at https://github.com/pachterlab/poseidon.

## Conclusion

Success and adoption of open source software demonstrates that reliable and powerful technologies can be realized through community development and improvement. We have developed a modular, highly customizable syringe pump array and USB microscope system, poseidon, with potential for broad application across the biological and chemical sciences. Syringe pumps can be used to operate microfluidic chips, control the chemical environment of a bioreactor, purify proteins, precisely add reagents to chemical reactions over time, or dispense specific amounts of fluid for any number of applications that require precise control of fluid flow. We have benchmarked the system against a commercial alternative in a demanding application: microfluidic emulsion generation. During design and testing of this system, we realized that our efforts would be wasted if new users could not rapidly implement and improve on our work. In developing the poseidon system as an open source hardware device we have illustrated six design principles that we hope can facilitate successful development of open source hardware devices (Fig. 5.)

## Acknowledgements

We thank Nicolas Bray and Kersh Theva for testing prototypes of the poseidon system and for valuable feedback. Thanks to Shannon Hateley for initial help with 3D printing and Zaid Adel Zayyad for help with Fig. 5.

## Author contributions statement

J.G. conceived of the project and developed the initial design for the syringe pumps. A.S.B. designed the syringe pump system and microscope, and implemented the poseidon software. E.V.B. helped with the design the poseidon system and oversaw hardware printing and design. A.S.B. and E.V.B. tested the poseidon system. J.G., A.S.B. and E.V.B. formulated the design principles. D.B. developed an initial version of the software. A.S.B., E.V.B., J.G. and L.P. wrote the manuscript.

## Supplementary Information

**Table 1.**
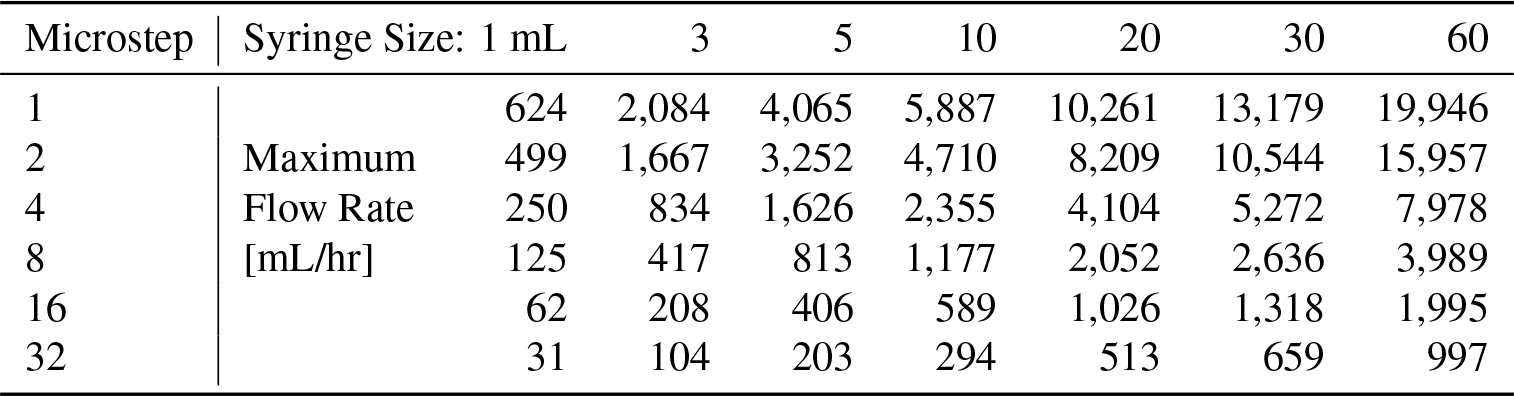
Maximum rates for given syringe and microstepping: Maximum flow rates [mL/hr] for a single pump for a given microstepping and Becton Dickinson (BD) syringe size. At lower flow rates, higher microstepping is desirable for a smoother flow.

**Table 2.** Stepper motor performance: Precision was calculated based on the steps per revolution and the pitch of the lead screw (0.8 mm/revolution). The maximum speed was measured for the displacement of the carriage with an error of 5.5% ± 1.5% of the set flow rate.

| Microstep | Steps per Revolution | Precision [$\mu\text{m}$] | Maximum Speed [mm/s] |
| --- | --- | --- | --- |
| 1 | 200 | 4 | 10 |
| 2 | 400 | 2 | 8 |
| 4 | 800 | 1 | 4 |
| 8 | 1600 | 0.5 | 2 |
| 16 | 3200 | 0.25 | 1 |
| 32 | 6400 | 0.125 | 0.5 |

## References

1. Gibb, A. Building Open Source Hardware: DIY Manufacturing for Hackers and Makers (Addison-Wesley Professional, 2014), 1 edn.

2. Arduino project. https://www.arduino.cc. Accessed: 2019-01-05.

3. Raspberry Pi foundation. https://www.raspberrypi.org. Accessed: 2018-10-06.

4. Pearce, J. M. Materials science. building research equipment with free, open-source hardware. Science 337, 1303–1304, DOI: 10.1126/science.1228183 (2012).

5. Pearce, J. Open-Source Lab: How to Build Your Own Hardware and Reduce Research Costs (Elsevier, 2014), 1 edn.

6. Maia Chagas, A. Haves and have nots must find a better way: The case for open scientific hardware. PLoS Biol 16, e3000014, DOI: 10.1371/journal.pbio.3000014 (2018).

7. Wijnen, B., Hunt, E. J., Anzalone, G. C. & Pearce, J. M. Open-source syringe pump library. PLoS ONE 9, e107216, DOI: 10.1371/journal.pone.0107216 (2014).

8. Maia Chagas, A., Prieto-Godino, L. L., Arrenberg, A. B. & Baden, T. A 3D-printable open-source platform for fluorescence microscopy, optogenetics, and accurate temperature control during behaviour of Zebrafish, Drosophila, and Caenorhabditis elegans. PLoS Biol 15, e2002702, DOI: 10.1371/journal.pbio.2002702 (2017).

9. Nuñez, I. et al. Low cost and open source multi-fluorescence imaging system for teaching and research in biology and bioengineering. PLoS ONE 12, e0187163, DOI: 10.1371/journal.pone.0187163 (2017).

10. Forman, C. J. et al. Openspritzer: an open hardware pressure ejection system for reliably delivering picolitre volumes. Sci Rep 7, 2188, DOI: 10.1038/s41598-017-02301-2 (2017).

11. Stephenson, W. et al. Single-cell RNA-seq of rheumatoid arthritis synovial tissue using low-cost microfluidic instrumentation. Nat Commun 9, 791, DOI: 10.1038/s41467-017-02659-x (2018).

12. Dolgin, E. How to start a lab when funds are tight. Nature 559, 291–293, DOI: 10.1038/d41586-018-05655-3 (2018).

13. Opentrons Inc. https://opentrons.com/ (2018). Accessed: 2018-11-30.

14. Wayland, M. T. & Landgraf, M. A cartesian coordinate robot for dispensing fruit fly food. J. Open Hardw. 2, 3, DOI: 10.5334/joh.9 (2018).

15. Carvalho, M. C. & Murray, R. H. Osmar, the open-source microsyringe autosampler. HardwareX 3, 10–38, DOI: 10.1016/j.ohx.2018.01.001 (2018).

16. Carvalho, M. C., Sanders, C. J. & Holloway, C. Auto-HPGe, an autosampler for gamma-ray spectroscopy using high-purity germanium (HPGe) detectors and heavy shields. HardwareX 4, e00040, DOI: 10.1016/j.ohx.2018.e00040 (2018).

17. Banović, L. & Vihar, B. Development of an extruder for open source 3D bioprinting. J. Open Hardw. 2, DOI: 10.5334/joh.6 (2018).

18. Fitzsimmons, R. E. et al. Generating vascular channels within hydrogel constructs using an economical open-source 3D bioprinter and thermoreversible gels. Bioprinting 9, 7–18, DOI: 10.1016/j.bprint.2018.02.001 (2018).

19. Macosko, E. Z. et al. Highly parallel genome-wide expression profiling of individual cells using nanoliter droplets. Cell 161, 1202–1214, DOI: 10.1016/j.cell.2015.05.002 (2015).

20. Zilionis, R. et al. Single-cell barcoding and sequencing using droplet microfluidics. Nat Protoc 12, 44–73, DOI: 10.1038/nprot.2016.154 (2017).

21. The Qt Company. https://doc.qt.io/qt-5 (2018). Accessed: 2019-01-03.

22. Autodesk fusion 360. https://www.autodesk.com/products/fusion-360. Accessed: 2018-10-07.

23. Harvard Apparatus. https://www.harvardapparatus.com/pumps-liquid-handling/syringe-pumps/infuse-withdraw/standard-infuse-withdraw-phd-ultra-syringe-pumps.html. Accessed: 2019-01-14.

